# A 4,000-year genomic history of Bantu speakers from the Congo rainforest

**DOI:** 10.64898/2026.10.27.762084

**Authors:** Oguzhan Parasayan, Maguelonne Roux, Fanny Mendisco, Javier Prado-Martinez, Dang Liu, Guillaume Laval, Christine Harmant, Marie-Hélène Pemonge, Cristina Pomilla, Liz Young, Matthew Hurles, Manj Sandhu, Diyendo Massilani, Bridget Chak, Luis B. Barreiro, Jean-Marie Hombert, Patrick Mouguiama-Daouda, Sacha Kacki, Rebecca Grollemund, Marie-France Deguilloux, Geoffroy de Saulieu, Richard Oslisly, Lluis Quintana-Murci, Etienne Patin

## Abstract

The Congo rainforest was the heartland of the Bantu Expansion, yet its demographic history remains partially understood. We inferred past population dynamics using genomes from 960 sub-Saharan Africans, including four newly sequenced ancient individuals from Iroungou cave and 176 present-day individuals from Gabon. Bantu-speaking populations began differentiating ∼4,000 years ago, preceding explosive population growth by a millennium. Distinct southward dispersals contributed to the ancestry of Bantu speakers in Gabon and elsewhere in sub-Saharan Africa, with the ∼700-year-old Iroungou individuals providing the closest genetic proxies for the latter. Both Bantu speakers and hunter-gatherers from the Congo rainforest experienced a major demographic crisis ∼1,500 years ago, coinciding with an archaeological hiatus. These findings illuminate early Bantu dispersals and the subsequent, unexplained collapse of rainforest populations.

## Main text

Although the demographic processes associated with the emergence of agriculture in Europe (*1*), East Asia (*2*) and Oceania (*3*) are now well understood, their reconstruction in sub-Saharan Africa has been hampered by the relative scarcity of archaeological records. Material-cultural and archaeobotanical data support the widespread emergence of village settlements and plant cultivation in Cameroon ∼3.0 thousand years ago (kya) (*4*) and a marked increase in human activity and iron working across the broader Congo rainforest ∼2.5 kya (*5–8*), including Cameroon, Gabon, Republic of Congo, Central African Republic and the Democratic Republic of Congo. In parallel, historical linguistics has shown that the earliest-diverging lineages of the Narrow Bantu language family, now widely spoken across sub-Saharan Africa, are found in present-day Cameroon (*9, 10*), supporting the hypothesis that Bantu-speaking populations (BSP) initially dispersed from this region before spreading across the subcontinent. Consistent with this model, genetic studies have shown that continent-wide BSP share recent ancestry with West-Central Africans(*11, 12*) and exhibit decreasing genetic diversity with increasing distance from the proposed Bantu homeland in Cameroon (*13*). Collectively, archaeological, linguistic and genetic findings support a model of the Bantu Expansion as a sequence of long-range dispersals of BSP from the Congo rainforest across sub-Saharan Africa, with the spread of agriculture and ironworking associated with later phases of this process.

Yet, despite broad concordance, several fundamental questions remain about the demographic processes underlying the early Bantu Expansion (*14*). First, when did the Bantu Expansion start? While Bantu-speaking populations are thought to emerge in Cameroon ∼5.0–3.0 kya (named here phase I, Fig. 1E) (*9, 10*), a widespread increase in human activity only started 2.5 kya (phase II, Fig. 1E) (*6*), raising the possibility that BSP diversification preceded demographic growth. Second, did the early Bantu Expansion involve sequential or parallel waves of migration? Whereas linguistic (*9, 10*) and genetic (*12, 13, 15, 16*) data for present-day BSP suggest an initial southward dispersal through a single interior route, the archaeological record supports two distinct movements along coastal and inland corridors (*4, 14*). Third, was occupation of the Congo rainforest by BSP continuous? Extensive field research and integrative analyses of radiocarbon dates have revealed a marked ‘hiatus’ in ceramic production and agricultural activities beginning ∼1.4 kya and lasting 600 years (phase III, Fig. 1E) (*7, 8, 17, 18*), reflecting either population collapse or changes in settlement patterns. Finally, why was admixture with local hunter-gatherers delayed in the Congo rainforest? Whereas admixture between incoming BSP and local populations in eastern and southern Africa occurred within a few centuries of their arrival, admixture with rainforest hunter-gatherers (RHG) begun more than a millennium after the initial Bantu settlement (*12, 13*). Here, we address these gaps by analyzing new whole genomes from ancient and present-day individuals in Gabon, a large forested region south of the Bantu homeland and located at the crossroads of the initial phase of Bantu migrations. By integrating genomic, archaeological and linguistic evidence, we reconstruct the early population dynamics, successive migrations, and demographic transformations that accompanied the spread of Bantu-speaking populations across sub-Saharan Africa.

**Fig. 1.**
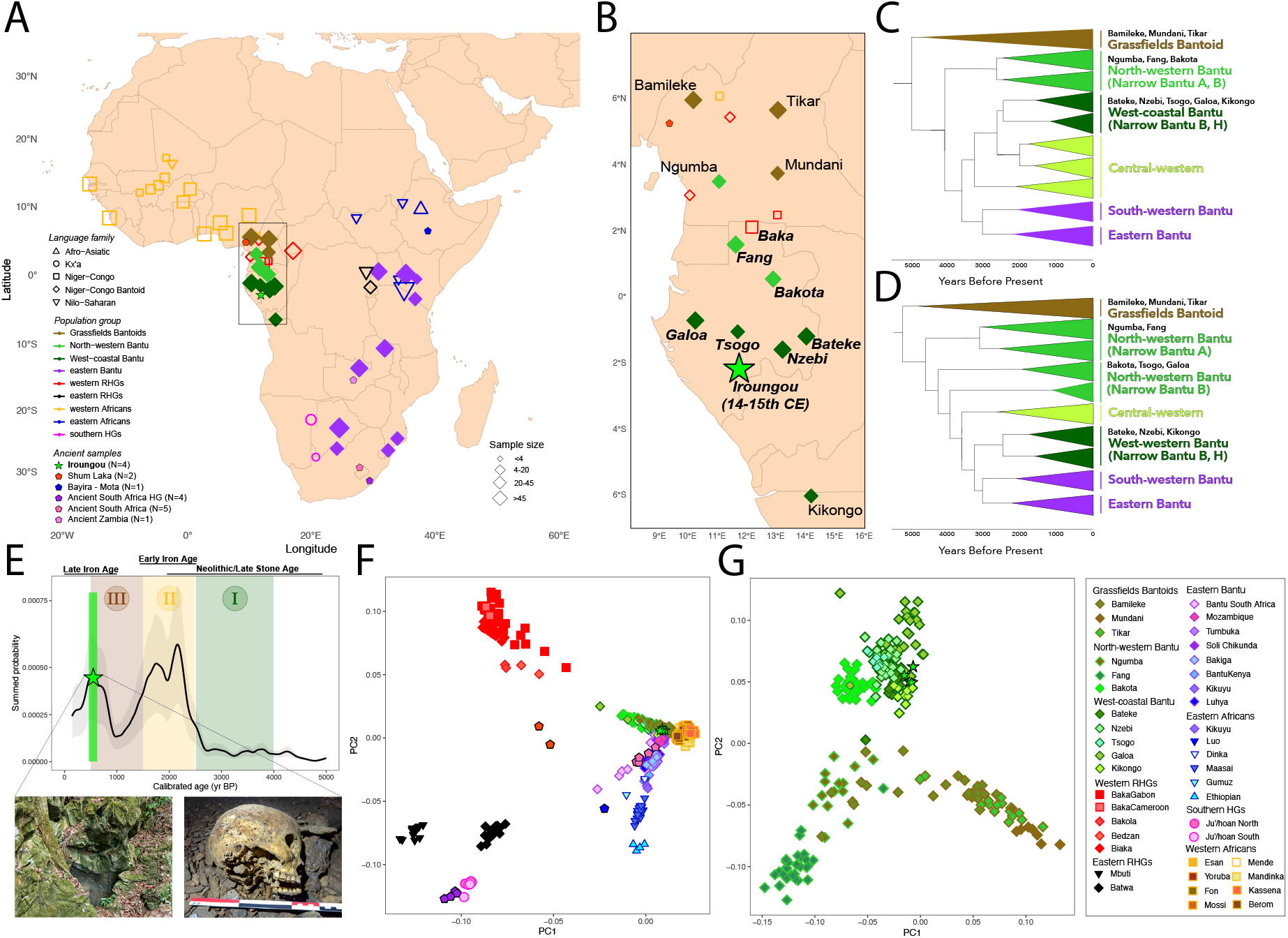
Genomic affinities among ancient and modern Bantu speakers. (A, B) Geographic location of sampled populations across (A) sub-Saharan Africa and (B) the Congo rainforest. The position, shape, color and size of points indicate the location, language family, source study and sample size of studied African populations, respectively. Outlined points indicate ancient individuals. Population names in italics indicate newly sequenced genomes generated in this study. (C, D) Simplified linguistic phylogeny of Bantoid languages spoken by the sampled populations, adapted from (C) Koile *et al.* (*10*) or (D) Grollemund *et al.* (*9*). (E) Summed probability distribution of radiocarbon dates in West-Central Africa, adapted from de Saulieu *et al.* (*7*). Colored areas indicate different phases of human activity; a low-activity stationary phase 4.0 – 2.5 kya overlapping the Neolithic (Phase I); a rapid growth associated with early Iron age 2.5 – 1.5 kya (Phase II); the archaeological hiatus 1.4 – 0.8 kya (Phase III). Green vertical lines indicate estimated dates of Iroungou ancient individuals (*22*). Photographs of (left) the entry of Iroungou burial cave and (right) a skull found at Iroungou, with typical pre-mortem incisive ablation. (F) Principal component analysis (PCA) of sub-Saharan Africans included in this study. (G) ChromoPainter-based PCA of Bantoid speakers from Westen Central Africa (*28*).

### Fine-scale genetic structure in the Congo rainforest

We generated high-coverage whole-genome sequences for 176 present-day individuals from Gabon (mean depth of coverage (DoC): 41×; Fig. 1A), including 146 sedentary villagers speaking North-Western or West-Coastal Bantu (*n*_pop_ = 6 populations; Fig. 1B–C; Table S1) – the most diverse early-diverging clades of the Narrow Bantu linguistic phylogeny (*9, 10*), and 30 Baka RHG. We jointly analyzed these genomes with available genome sequences from 768 sub-Saharan Africans and 80 non-Africans, encompassing populations speaking Grassfields languages (*n*_pop_ = 3) – the Non-Bantu languages most closely related to Narrow Bantu –, and Eastern Bantu (*n*_pop_ = 10) languages (Fig. 1B–C). The dataset also included RHG, Kx’a-speaking Ju’/hoan, and Nilo-Saharan-speaking populations, representing proxies of sources contributing non-Bantu-related ancestry in BSP from west-central, southern and eastern Africa, respectively (*15, 16, 19–21*). After quality-control filtering (Fig. S1), the resulting dataset comprised 1,024 individuals from 50 populations, and 56,787,614 biallelic SNPs.

In addition, we screened 22 ancient individuals from the Iroungou burial cave in Gabon, dated to 0.6–0.7 kya (Fig. 1D,E) (*22*), for endogenous DNA. Intriguingly, all adult individuals at Iroungou exhibit pre-mortem ablation of the four upper incisors, a cultural practice no longer observed among BSP today. We obtained whole-genome sequences for four individuals at DoC ranging from 0.9× to 7.2× (Table S2). These genomes were imputed, together with other available ancient African genomes, using our modern genome dataset as a reference panel (*23*). To assess imputation accuracy, we down-sampled sequencing reads from four deeply sequenced ancient genomes (DoC >10×), including a Bantu-related individual from Eland Cave (*24*). When restricting to variants with minor allele frequency (MAF) ≥1%, imputation accuracy (*r*²) exceeded 0.95 at DoC ≥0.5× (Fig. S2, Table S3). We therefore applied these filters to the observed ancient data.

We then investigated genetic affinities among BSP, Iroungou individuals, and other African populations (*23*). Principal Component Analysis (PCA) (*25*) and ADMIXTURE (*26*) analyses supported close affinities between BSP from west-central, eastern, and southern Africa and Grassfields speakers from Cameroon, together with weaker affinities to RHG, Kx’a-speaking Ju’/hoan and Nilo-Saharan speakers from regions where BSP respectfully expanded (Fig. 1F; Fig. S3). These results are consistent with previous studies reporting a west-central African origin for BSP and subsequent admixture with local populations during their expansion (*11–13, 15, 16, 20, 27*). Haplotype-based PCA (*28*) revealed fine-scale structure among Bantu-related populations from the Congo rainforest, with individuals broadly clustering according to Grassfields, North-Western Bantu, and West-Coastal Bantu linguistic affiliations (Fig. 1G, Fig. S4). The Iroungou individuals clustered most closely with present-day West-Coastal BSP, particularly the Kikongo from Angola (Fig. S5), supporting their affiliation with BSP. Together, these results establish a genomic framework for investigating demographic changes associated with the emergence of BSP in the Congo rainforest and subsequent contribution to continent-wide dispersals.

### Differentiation among Bantu-speaking populations preceded demographic growth

The Bantu Expansion is thought to have begun ∼5.0–4.0 kya with the diversification of small Bantu-speaking communities (*9, 29*). However, previous estimates of population size changes and divergence times among BSP have been inconsistent (*8, 16, 30*), due to the use of ascertained data and low-resolution or biased approaches. We inferred temporal changes in the effective population size (*N*_e_) of BSP using CoalRate (*31*), which estimates ancestry-specific coalescence rates by combining genome-wide genealogies (*32*) with local ancestry inference (*33*) to account for biases introduced by admixture (*34*). We confirmed with simulations that CoalRate can detect expansions and bottlenecks occurring over the past five millennia (Fig. S6). We found that all Niger-Congo-speaking populations – the broader language family including Bantu languages – shared similar *N*_e_ trajectories until approximately 10 kya (Fig. 2A; Table S4), when Grassfields- and Bantu-speaking groups separated from West African populations, including Yoruba and Esan from Nigeria. Strikingly, between 4.0 and 2.5 kya, coinciding with phase I (Fig. 1E), *N*_e_ of all BSP reduced to ∼4,000, while *N*_e_ in related West African populations increased to ∼10,000. A low *N*_e_ was also observed among Grassfields speakers (i.e., Tikar from Cameroon), suggesting that this phenomenon preceded the emergence of Narrow Bantu languages. Following this contraction, BSP exhibited sharp population growth starting 2.5 kya (Fig. 2A), coinciding with archaeological evidence for increased human activity (phase II, Fig. 1E) (*5–8*).

**Fig. 2.**
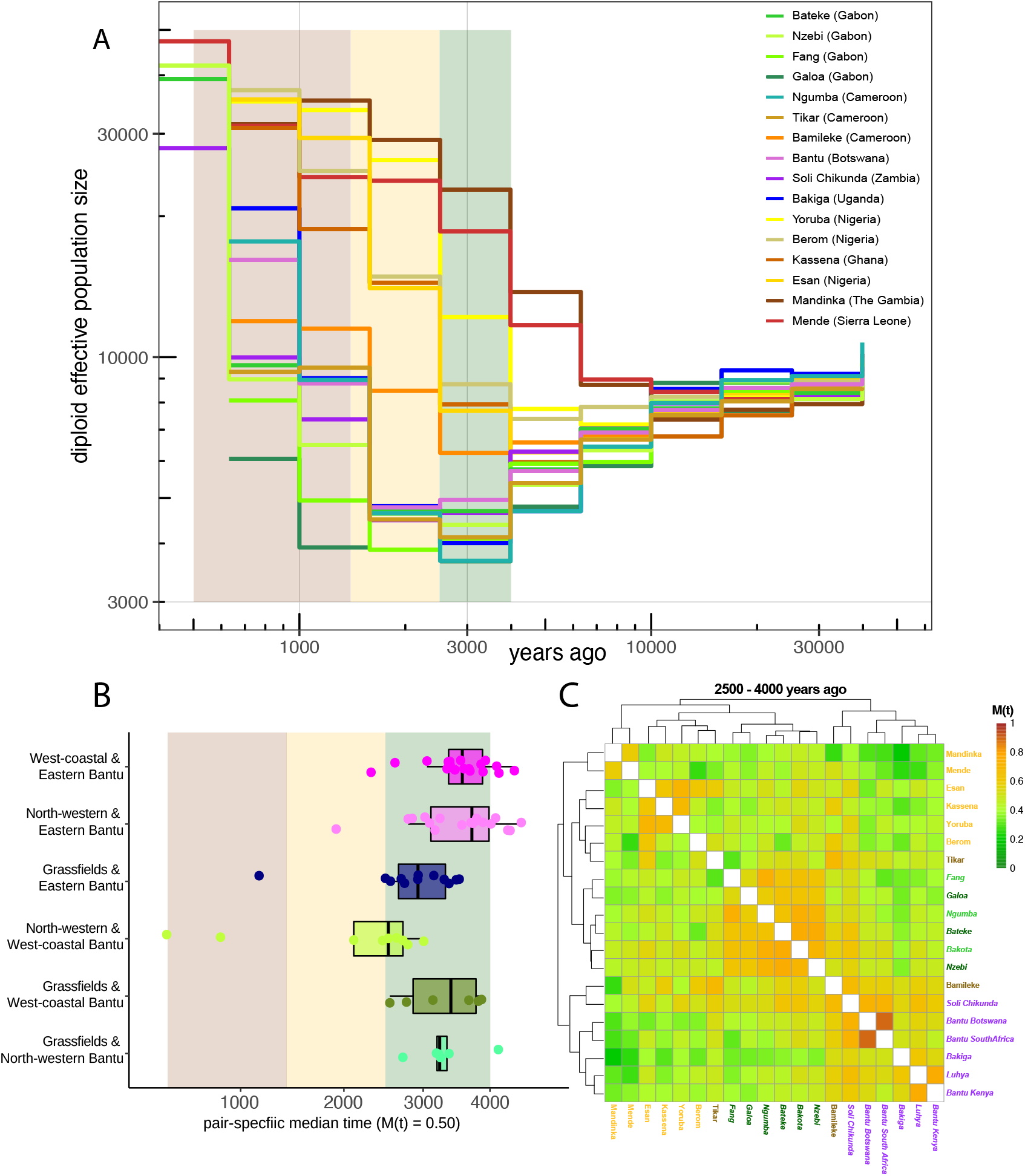
Effective population sizes and divergence times. (A) Effective population size (*N*_e_) trajectories for western and west-central African populations, estimated with CoalRate (*31*). *N*_e_ (y-axis) and time (x-axis) are on log-scale. Colored areas indicate high and low activity periods, as in Fig. 1E. (B) Distribution of pairwise population split times (years ago) estimated using *MSMC-IM* (*35*). Each point corresponds to the divergence time – defined as the time point by which 50% of migration events between the two populations occurred – between a pair of populations speaking languages indicated on the y-axis. (C) Heatmap of pairwise cross-population *M*(*t*) values averaged over Phase I (4.0–2.5 kya). Population name colors correspond to linguistic groups in Fig. 1C,D. Population names in italics indicate groups whose non-Bantu ancestries are masked.

The low *N*_e_ detected during Phase I can result from a founder event in the ancestral population of Bantu speakers and/or subdivision into small, isolated demes. To distinguish between these scenarios, we estimated divergence times among BSP by fitting an Isolation-with-Migration (IM) model with *MSMC-IM* (*35*) to ancestry-specific cross-coalescence rates (*23*). In line with previous estimates (*16*), most BSP diverged between 4.0 and 2.5 kya (Figs. 2B,C and S7), supporting the view that their reduced *N*_e_ coincided with population subdivision. Collectively, these findings suggest that the Bantu Expansion occurred in two broad demographic phases: an early subdivision phase between 4.0 and 2.5 kya, characterized by low population sizes and coinciding with limited human activity (phase I, Fig. 1E), followed from 2.5 kya onward by rapid demographic growth and regional diversification (phase II, Fig. 1E).

### Southward Bantu dispersals followed distinct coastal and inland routes

Previous genetic studies have shown that Eastern BSP share greater genetic affinities with present-day populations from southern than northern Cameroon, a pattern that has been interpreted as evidence for an initial southward direction of Bantu dispersals (*12, 13, 15, 16*). We used our dense modern and ancient whole-genome dataset to investigate whether these Bantu southward dispersals proceeded along a single route through the Congo rainforest, as suggested by linguistic phylogenies (*9, 10*), or along two routes, following the Atlantic coast and an interior corridor toward the Congo Basin, as suggested by the location of major Neolithic archaeological sites (*4, 14*). Because the archaeological record places the proposed coastal dispersal through Gabon, we tested these scenarios by examining whether Gabonese BSP and BSP elsewhere in sub-Saharan Africa derive Bantu-related ancestry from the same source or different sources. Specifically, under the single-dispersal scenario, both Gabonese and Eastern BSP are expected to derive Bantu-related ancestry from Grassfields-speaking populations in Cameroon, whereas the two-dispersal model predicts that Eastern and Gabonese BSP derive ancestry from Grassfields-speaking populations and North-Western BSP from Cameroon (hypothesized to have been present along the Atlantic coast preceding West-Coastal BSP (*23, 36*)), respectively.

We used SOURCEFIND (*37*) to decompose Bantu-related ancestry into Grassfields- and North-Western Bantu-related components, corresponding to the two major genetic clusters identified in Cameroon (Fig. 1G). The sources for these components included Bamileke from western Cameroon, as previously observed (*38*), and Ngumba from southern Cameroon, respectively. All present-day Eastern BSP showed 40−60% of each component (Fig. 3A; Table S5), suggesting that they descend from the same ancestral population. In contrast, North-Western Bantu-related ancestry accounted for 83−99% of Bantu-related ancestry in Gabonese BSP, indicating closer relatedness to southern than to northern Cameroonian populations, consistent with the proposed coastal route through Gabon.

**Fig. 3.**
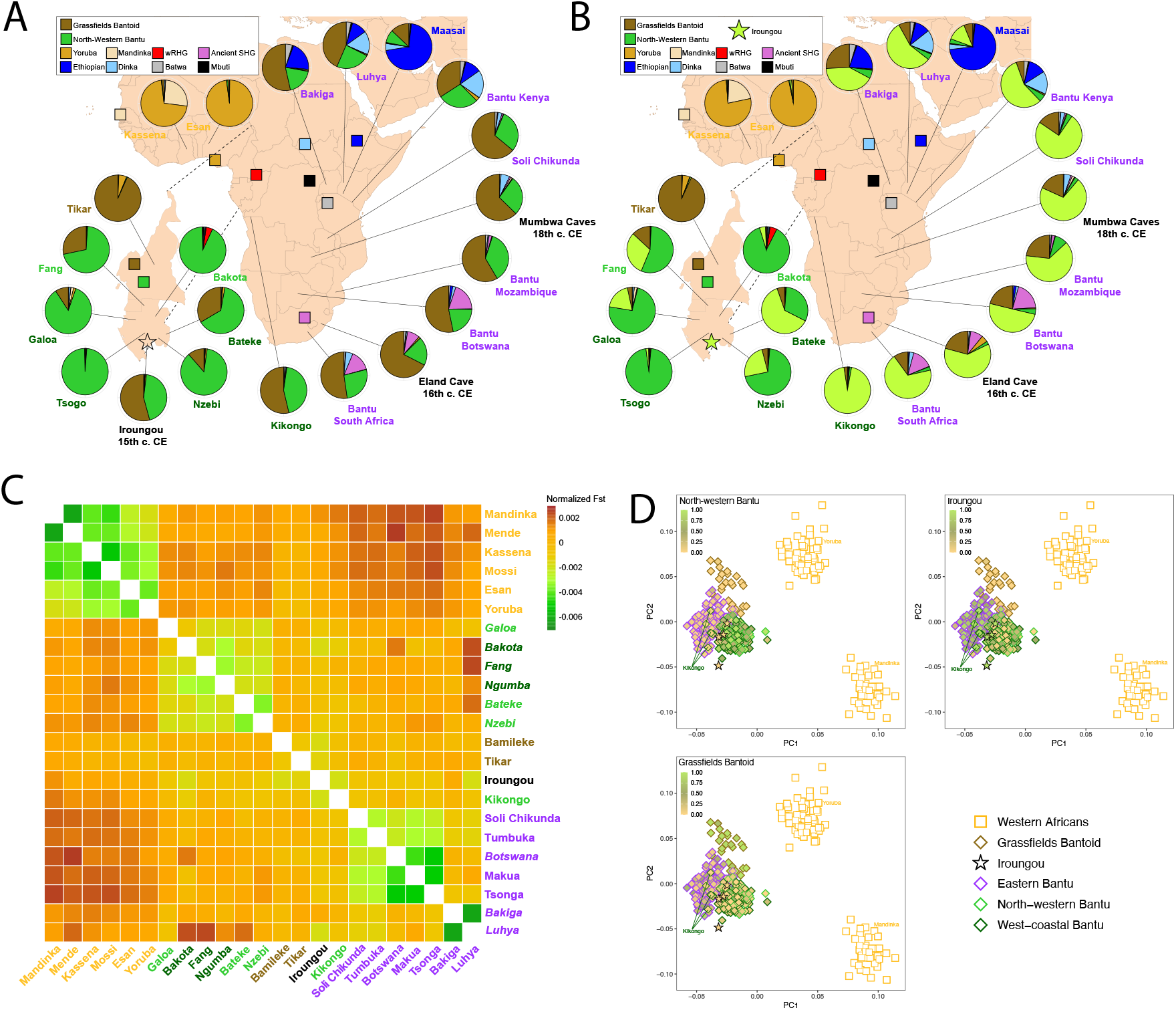
Sources of Bantu-related ancestry. (A) Genetic contributions of Grassfields- and North Western Bantu-speaking populations from Cameroon, together with other non-Bantu-related sources, in BSP, estimated with SOURCEFIND (*37*). (B) Genetic contributions of Iroungou ancient individuals from Gabon, Grassfields- and North Western Bantu-speaking populations from Cameroon, together with other non-Bantu-related sources, in BSP. (C) Heatmap of hierarchical clustering of pairwise Hudson *F*_ST_ normalized using double-centering. Population name colors correspond to linguistic groups in Fig. 1C,D. Population names in italics indicate groups whose non-Bantu ancestries are masked. (D) Ancestry-specific PCA of Bantu-related ancestry in BSP, colored according to proportions of North-Western Bantu-, Iroungou-, and Grassfields-related components.

To resolve further the ancestry associated with the putative interior route, we searched for BSP from west-central Africa that represent the closest source for Bantu-related ancestry in Eastern BSP. We found that ancient Iroungou individuals were the best-matching source, when compared to their closest present-day relatives (Fig. S8), the Kikongo from Angola (Table S6), who have been identified as the closest source among present-day BSP (*12, 13, 15, 16*). Considering Grassfields-speaking population, North-Western BSP, and Iroungou individuals (*n* = 4) as sources in SOURCEFIND, Iroungou individuals contributed the most Bantu-related ancestry in Eastern BSP (56−84%; Fig. 3B), while contributing 2%−30% in present-day Gabonese BSP, with the notable exception of Bateke (63%; Fig. 3B). Consistent with these results, PCA and *F*_ST_ statistics computed after non-Bantu-related ancestry masking indicated that Eastern BSP shared greater affinities with Iroungou, and Grassfields-speaking individuals (Fig. 3C,D), whereas West-Coastal BSP from Gabon broadly overlapped with North-Western Bantu speakers from Cameroon, in contrast with West-Coastal BSP from Angola. Furthermore, we verified that the genetic affinities observed among BSP were not explained by an unmodeled source of Bantu-related ancestry, carried by Ubangi-speaking populations from Central African Republic (*39*), and found that Ubangi-related component was not correlated with any of the three components studied here (Fig. S9).

To evaluate the timing of these dispersals, we examined time-dependent migration probabilities *M*(*t*), which indicated that ancestors of Grassfields speakers and Eastern BSP separated from ancestors of North-Western and West-Coastal BSP during Phase I (4.0 – 2.5 kya), consistent with the early formation of distinct inland and coastal lineages (Fig. 2B). *M*(*t*) distributions further supported a split between North-Western and West-Coastal BSP at the beginning of Phase II, coinciding with the earliest evidence for possible Bantu-associated occupation of coastal Gabon (Fig. 2B; Fig. S7) (*4*). These results corroborate the coastal route model and demonstrate that more recent population movements from Cameroon into Gabon, such as those of Fang speakers during 18–19th century (*40*), cannot explain the genetic patterns observed. Collectively, these findings support the view that BSP derive from distinct dispersal events, with the coastal and inland routes carrying ancestry related to BSP from southern Cameroon and northern Angola/ancient southern Gabon, respectively.

### Congo rainforest populations collapsed ∼1.4 kya

Radiocarbon-dated archaeological records from the Congo rainforest have provided evidence for a hiatus in human activity, which has been interpreted as a widespread demographic collapse between 1.4 and 0.8 kya (*5, 7, 8*). To support this hypothesis, Seidensticker *et al*. inferred how *N*_e_ has changed through time in Gabonese BSP, using IBDNe (*41*). They concluded that *N*_e_ remained low over the past 2,500 years before rapidly increasing ∼0.5 kya (*8*). They interpreted these results as evidence for a ‘spread-over-spread’ model, in which BSP that first expanded in Gabon and underwent the demographic collapse were replaced by present-day Gabonese BSP, since the latter do no show genomic signatures of either event. However, this interpretation has been questioned because IBDNe fails to capture such successive demographic events (*38*).

To assess whether the hiatus could leave detectable genomic signatures, we simulated demographic models incorporating realistic combinations of growth and collapse events over the past 3,500 years (*23*). We compared true simulated *N*_e_ values with estimates obtained using CoalRate (*31*) and HapNe-IBD, a new method that controls for spurious *N*_e_ oscillations in recent generations and outperforms IBDNe (*42*). HapNe-IBD better captured growth-decline-growth models occurring over the past 3,000 years, compared to CoalRate (Figs. 4A and S10). We therefore applied HapNe-IBD to the empirical data. In Gabonese BSP, *N*_e_ increased rapidly from 2.5 kya onward, coinciding closely with the intensified human activity documented in the archaeological record (phase II, Fig. 1E) (*7, 8, 17, 18*). Strikingly, this growth was interrupted by an abrupt *N*_e_ decline ∼1.4 kya (Fig. 4B), providing independent genetic evidence for a major demographic contraction coinciding with the archaeological hiatus (phase III, Fig. 1E).

**Fig. 4.**
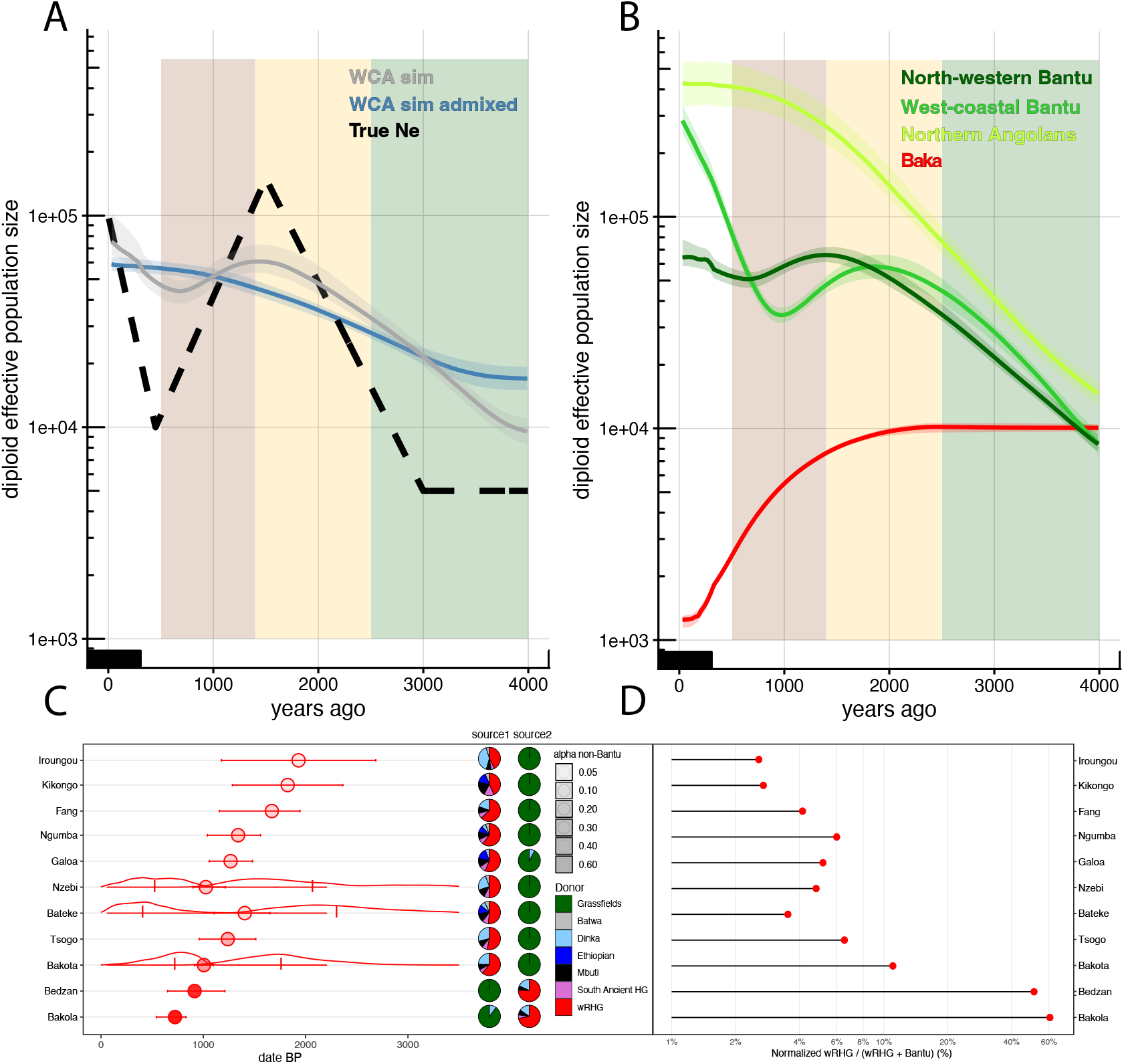
Demographic changes and admixture during the hiatus. (A) Estimated *N*_e_ trajectories for a simulated population undergoing a 10-fold size reduction during the hiatus, using HapNe-IBD (*42*). The black, grey and blue lines indicate the simulated model, and HapNe-IBD estimates on the simulated data in absence or presence of admixture from a third population, respectively. Colored areas indicate high and low activity periods, as in Fig. 1E. (B) Estimated *N*_e_ trajectories for Gabonese and Angolan BSP, using HapNe-IBD. (C) Estimated times and sources for admixture events between BSP and RHG, using fastGLOBETROTTER (*44*). Points and bars indicate the point estimate and confidence interval of the admixture time under a single-pulse model. The distributions indicate posterior distributions of the admixture time under a two-pulse model. Pies indicate the relative contributions of source populations in the two admixing sources. (D) Normalized RHG-related ancestry proportions estimated using SOURCEFIND (*37*) for the same populations shown in Fig. 4C.

Several explanations have been proposed for this demographic collapse, including long-term environmental changes affecting subsistence and settlement patterns, and a prolonged epidemic (*7, 8, 17, 18*). We reasoned that hunter-gatherers from the Congo rainforest (RHG) might have been less vulnerable to disruptions in agricultural production, given their diverse strategies for subsisting in the Congo rainforest (*43*), whereas an infectious disease outbreak could have affected both groups. We detected an abrupt decline in *N*_e_ ∼1.4 kya in Baka RHG, comparable in timing and magnitude to that inferred in Gabonese BSP (Fig. 4B). This shared demographic contraction is consistent with a crisis affecting both farming and hunter-gatherer populations.

We next investigated whether admixture between BSP and RHG preceded or followed the demographic contraction, as interactions between these populations may have facilitated pathogen transmission. We estimated the timing and sources of admixture between the BSP and RHG, using fastGLOBETROTTER (*44*). Consistent with previous findings (*12, 30*), we detected reciprocal admixture in all tested populations, using minimally admixed Grassfields speakers and Baka RHG as sources and 1,000 bootstrap iterations (*23*). Co-ancestry decay curves supported either a single-pulse admixture event occurring > 1.5 kya or a two-pulse event starting ∼2.0 kya in 6 out of 9 BSP (Fig. 4C), broadly preceding the archaeological hiatus (*7, 8, 17, 18*). Together, these findings reveal a major demographic contraction affecting both ancestors of present-day Bantu-speaking agriculturalists and rainforest hunter-gatherers, consistent with a widespread crisis for which epidemic disease remains a plausible, although unconfirmed, explanation.

## Discussion

Despite major advances in archaeology and historical linguistics (*45*), the demographic processes leading to the emergence of Bantu-speaking populations and the spread of village settlements, iron working, agriculture and Bantu languages have remained partially understood. Here, we assembled a large genomic dataset of BSP, including new genomes from ancient and modern Gabonese, to reconstruct the early stages of the Bantu Expansion. We show that BSP started to differentiate between 4.0 and 2.5 kya, coinciding with Phase I, a period of low human activity and effective size, followed by sustained demographic growth starting ∼2.5 kya. This finding supports the view that early Bantu dispersals within the Congo rainforest predated the spread of metallurgy and agriculture.

From ∼3,000 years onward, ceramic traditions diversified into at least two major streams: the Obobogo- and the Imbonga-related cultures, which flourished along the Atlantic coast of Cameroon and Gabon and in the inner Congo basin, respectively (*4*). Conversely, linguistic phylogenies support a single interior route through the Congo rainforest (*9, 10*), suggesting that the languages once spoken by the populations that took the coastal route vanished (*36*). Consistent with the archaeological data, our analyses reveal that Gabon was likely settled by BSP related to North-Western Bantu speakers, through a distinct dispersal than the rest of sub-Saharan Africa. Gabonese BSP might have later acquired West-Coastal Bantu languages (*36*), which could explain why no linguistic clade reflects the coastal diffusion of Obobogo-associated cultures. Notably, despite speaking the same language family, present-day West-Coastal BSP from Gabon and Angola carry largely different proportions of the North-Western Bantu-related component, consistent with the occurrence of a language shift. This scenario is also consistent with the unusual linguistic features in some West-Coastal Bantu languages that may be explained by retention of extinct North-Western Bantu strata (*46, 47*).

Although Gabonese BSP derive most of their ancestry from the proposed coastal dispersal, some populations in southern Gabon carry ancestry from the inland migration, most notably the Bateke, suggesting the Bateke Plateau and the Ogooué as plausible contact zone between descendants of the two population movements. Interestingly, the earliest dated smelting in Gabon is at La Lopé (∼2.4 kya) (*48*), contemporaneous with Imbonga-related settlement of the inner Congo basin (*4*). However, older smelting evidence lies further north, including in southern Cameroon (∼2.6 kya), so the coastal and inland populations may both have obtained metallurgy from a common northern source.

Our results also identify the ∼600-year old individuals from the Iroungou cave (Gabon) as the closest available genetic proxy for Bantu-related ancestry in virtually all BSP inhabiting outside the Congo rainforest, confirming a southward route for the Bantu Expansion (*12, 13, 15, 16*). Iroungou individuals were genetically closer to present-day BSP from Angola than to those currently inhabiting Gabon, suggesting a localized turnover. However, copper artefacts from Iroungou cave likely originated from the Democratic Republic of Congo, raising the possibility that these ancient individuals were local migrants (*22*). Deep sampling of present-day populations from the region will be needed to demonstrate local genetic discontinuity.

Finally, we provide independent genomic evidence for a major population collapse in the Congo rainforest ∼1.4 kya, corroborating the demographic interpretation of the archaeological hiatus (*7, 8, 17, 18*). The inferred contraction affected both Bantu-speaking agriculturalists and hunter-gatherers of the Congo rainforest, indicating that the crisis extended beyond communities dependent on iron working and cultivated plants. Several mechanisms could account for this widespread disruption, including climate-driven changes in resource availability and epidemics. The similar timing and magnitude of the demographic contractions inferred in both groups, together with the duration of the archaeological hiatus, are consistent with a prolonged epidemic crisis, as previously hypothesized (*17*). Nevertheless, shared demographic trajectories do not uniquely implicate infectious disease: environmental disturbances could have affected multiple subsistence systems, leading to the observed reductions in effective population size. Ancient pathogen genomics will be essential to distinguish among these scenarios and, if an epidemic occurred, identify the responsible pathogen.

## Conclusion

Our results reveal the Bantu Expansion as a complex, multi-stage demographic process rather than a single wave of population growth and dispersal. Population differentiation in the proposed Bantu homeland ∼4–3 kya preceded demographic growth and large-scale migrations, with distinct coastal and inland dispersal trajectories contributing differently to the ancestry of present-day Bantu-speaking populations across sub-Saharan Africa. Furthermore, genomic evidence for a major population collapse in the Congo rainforest ∼1.4 kya, affecting both Bantu-speaking agriculturalists and rainforest hunter-gatherers, reveals substantial demographic disruption following initial settlement. Time-stratified ancient genomes from across Central Africa will be essential to resolve population continuity, replacement, and mobility throughout these successive stages.

Materials and methods are available in the supplementary materials.

## Materials and Methods

### Ethical approval and informed consent

The study was conducted in accordance with applicable national regulations and international ethical principles governing human genetic research. Samples were originally collected in Gabon in 2003 under research authorization no. 007/MESRIT/SG/DRCST, issued by the Gabonese Directorate of Research and Scientific and Technical Cooperation on 2 July 2003. All participants provided written informed consent for blood collection and genetic research. The subsequent genomic research protocol (2011–54) received approval from the Institutional Review Board of Institut Pasteur (IRB00006966) on 6 January 2012, followed by annual ethical reviews. The retrospective Gabon-Div study (protocol 2015-121) received a favorable opinion from the National Ethics Committee of Gabon (PROT No. 0016/2016/SG/CNE), subject to the involvement of designated Gabonese scientific experts. Biological samples were anonymized, and no correspondence between participant identities and sample identifiers was retained.

### Participant recruitment and sample collection

Blood samples were collected in Gabon in 2003 as part of a collaborative survey designed to capture the country’s ethnolinguistic and genetic diversity. The original collection comprised 890 healthy adult male volunteers from 20 ethnolinguistically defined populations, including Bantu-speaking agriculturalists and Central African rainforest hunter-gatherers. Eligible participants were adults who consented to blood collection and genetic research. Recruitment and sampling took place in local dispensaries or facilities suitable for medical care near participating villages. Field teams included a researcher, a nurse and a Gabonese anthropology student, who documented linguistic and ethnolinguistic information, assessed reported kinship relationships and ensured appropriate blood collection procedures. Participants were informed of the minimal risks associated with blood sampling, and pharmaceutical products were offered as compensation for participation. Blood samples were transported to the Centre International de Recherches Médicales de Franceville (CIRMF), where DNA was extracted. DNA samples were coded and transferred to Institut Pasteur in Paris under the original Gabonese research authorization, and subsequently stored at −80°C for genomic analyses.

### Whole genome sequencing and processing for present-day individuals

DNA libraries for 176 eligible samples were prepared at the Wellcome Sanger Institute (Cambridge, UK), using the Illumina TruSeq DNA PCR-free Library Preparation Kits. All libraries were prepared according to the manufacturer’s instructions and were sequenced on the Illumina NovaSeq 6000 platform (Illumina Inc., USA) to generate 150-bp paired-end reads and obtain ∼30× depth of sequencing coverage per individual.

To minimize batch effects, both newly sequenced genomes and previously published genomes were subjected to the same data processing pipeline. Sequence quality of *FASTQ* files were assessed using the FastQC software (*49*) and multiQC v.1.9 (*50*) followed by conversion to unmapped *BAM* files (*uBAM*). Read groups were added, and Illumina adapters were marked using Picard tools v.2.20.1 (https://broadinstitute.github.io/picard). Read pairs were mapped to the hs37d5 human reference genome (*51*) with the Burrows-Wheeler Aligner (BWA mem) v.0.7.17 (*52*) and duplicates were marked with Picard. Per-sample depth of coverage (DP) and the number of reads mapped to the human reference genome were calculated using SAMtools v.1.10 (*53*). DNA contamination levels were estimated with VerifyBamID2 (*54*). Sequencing read mapping qualities were adjusted using capmq (https://github.com/mcshane/capmq) to account for estimated DNA contamination. Samples with a FREEMIX contamination estimate exceeding 1% were excluded, resulting in the removal of 13 sub-Saharan African samples (Table S1).

### Genotype calling, quality filters, and phasing

Variant calling was performed with GATK v.4.1.2.0, following GATK Best Practices (https://software.broadinstitute.org/gatk/best-practices/). Samples were processed independently with HaplotypeCaller, and *gVCF* files were merged using CombineGVCFs. Joint genotype calling across all samples was then performed using the GenotypeGVCFs tool with the option --allSites. Variant sites were extracted and then filtered using GATK variant quality score recalibration (VQSR) applying a truth sensitivity threshold of 99.5 for SNPs, yielding 56,787,614 SNPs on autosomes. Indels were excluded from downstream analyses. Genotypes at variant and invariant positions were set to missing if their sequencing depth fell outside the sample-specific mean ± standard deviation or their genotype quality (GQ/RGQ) was below 30, using BCFtools v1.11 (Li, 2011). For samples with more than 5% missing data (59), the minimum GQ/RGQ threshold was lowered to 10.

Cryptic relatedness and duplicate samples were identified by estimating kinship coefficients with KING v.2.2.7 (*55*), as implemented in PLINK2 (*56*). Prior to kinship analysis, variants with a genotyping rate below 90% were removed. Individuals presenting a kinship coefficient greater than 0.1 excluded until no related pairs remained. When ties occurred, the individual with the lowest sequencing depth was discarded. This filtering step led to the removal of 56 previously published genomes. Deviations from Hardy–Weinberg equilibrium were assessed using RUTH (*57*), which explicitly models population structure. The first ten principal components derived from the genotype data were included as covariates, and 8,196 variants with *P*-values below 10⁻³ were excluded. Subsequently, the dataset was then phased and imputed using SHAPEIT4 v.4.2.2 (*58*) using following parameters --mcmc-iterations 10b,1p,1b,1p,1b,1p,1b,1p,10m --pbwt-depth 8 –sequencing.

### Ancient DNA extraction and library construction

Ancient DNA extraction and library construction was carried out in the cleanroom facilities of the Laboratory PACEA, Palaeogenetic Platform, Bordeaux University, France. All teeth were decontaminated using a 3% bleach solution and sectioned along the cementum-enamel junction. Powdered material was obtained either by drilling into the pulp chamber or by grinding the entire tooth into a fine powder. Approximately 50 mg of powder was used for each DNA extraction. The extraction procedure followed the protocol published by Velsko et al. (*59*). Two libraries were prepared per individual using a partial uracil-DNA-glycosylase (UDG) double-stranded library preparation protocol with unique dual-index combinations, following Aron and Brandt (*60*). Sequencing was performed by the iGenSeq Genotyping and Sequencing Platform at the Paris Brain Institute (ICM) using an Illumina NovaSeq X Plus with a 10B Reagent Kit (200 cycles).

### Whole genome processing for ancient individuals

Sequence quality of *FASTQ* files were assessed using the FastQC software (*49*). Paired-end reads were merged and trimmed for adapters using leeHom (*61*). Reads larger than 28 base pairs were selected using awk and mapped onto the hs37d5 human reference genome (*51*) using BWA aln v.0.7.17 (*52*). Mapping parameters depended on the treatment: for non-UDG libraries, -t 4 -l 0 -n 0.01 was used, whereas for UDG-treated libraries, a seed length of 18 (-l 18) was specified. Aligned sequences in BAM format were processed with SAMtools v1.10 (*53*) to create index files and flagstat reports and to select and sort mapped reads. Re-sequenced libraries merged before the duplicate removal by SAMtools. Picard tools (http://broadinstitute.github.io/picard/) was used to remove duplicates. Multiple unique libraries were merged by before filtering by mapping quality (--q 20). Damage patterns were analyzed using mapDamage v.2.2.1 (*62*), followed by end-clipping using BamUtil (https://genome.sph.umich.edu/ wiki/BamUtil:_trimBam). According to the damage pattern, three to four bases at the ends of each read were clipped for the USER-treated libraries and 10 bases for non–USER-treated libraries.

### Imputation and imputation quality assessment

GLIMPSE (v.1.1.1) (*63*) was used to impute 14 ancient genomes including four Iroungou individuals (Table S2). We followed the workflow described in the online tutorial (https://odelaneau.github.io/GLIMPSE/glimpse1/tutorial_hg19.html), using Genotype Likelihoods (GL) computed with Bcftools (mpileup --I --E, call --Aim parameters) using the newly constructed high coverage Sub-Saharan African dataset as reference panel instead of 1000G reference panel phase 3. To assess the imputation accuracy we down-sampled four high coverage (>10x) ancient genomes (BaA001, Ela001, I10871, I5950) (*24, 64, 65*), which were included in the joint genotype calling together with high coverage present-day individuals, to the mean depth coverages at 0.3, 0.4, 0.5, 1, 2, 10x. After excluding four true ancient genomes from the reference panel, each down-sampled genome was separately imputed with following the GLIMPSE workflow. GLIMPSE_concordance was run on imputed and high-coverage genomes (--minDP 8 --bins 0 0.0001 0.0005 0.001 0.005 0.01 0.05 0.1 0.5) as described in the online tutorial. The variants with MAF <1% in the reference panel were filtered out and imputed genotypes with genotype posterior probabilities (GP) <0.90 were set as missing using BCFtools in the imputed dataset including Iroungou and published ancient genomes. Imputed and filtered ancient dataset was merged with high coverage Sub-Saharan African dataset.

### smartPCA and ADMIXTURE

The VCF files, including imputed ancient genomes, were converted to PLINK bed/bim/fam format and filtered to remove sites with missing genotypes or minor allele frequency (MAF) < 1% using PLINK v1.9. ADMIXTURE v1.3.0 (*26*) was run on the merged dataset with K values ranging from 2 to 12 using default parameters. Principal component analysis (PCA) was performed with smartPCA from EIGENSOFT v7.2.1 to calculate eigenvectors (*25*).

### Chromosome painting based approaches

#### Haplotype-based PCA and fineSTRUCTURE

We ran ChromoPainter/fineSTRUCTURE (*28*) on the imputed and phased data that are filtered for MAF<1%. After estimation of switch and mutation rates using 10 expectation-maximization (EM) iterations, each haplotype was painted using all others (-a 0 0) to generate pairwise coancestry matrices. All individuals were initially placed in a single cluster, and fineSTRUCTURE MCMC was run on the resulting coancestry matrices for 3 million iterations with a 2 million iteration burn-in, sampling every 100 iterations. The MCMC sample with the highest posterior probability was used as the starting point for 100,000 hill-climbing moves to reach the final inferred state. (Clustering: -s1 -c2.0 -x3M -y2M -z100 | Hill-climbing: -x0 -y100k -z100 | Tree-building: -mT -x0 -k2 -T1). The PCA and the dendrogram constructions were performed using the R functions from FinestructureLibrary.R (https://people.maths.bris.ac.uk/∼madjl/finestructure/finestructureR.html).

#### Admixture inference with SOURCEFIND and fastGLOBETROTTER

We ran ChromoPainter v2 (*28*) with 10 iterations of EM to infer parameters on chromosomes 1, 8, 15, and 22. We then generated the copy vector and painting sample files for the 22 autosomes using wRHG Mandinka, Mbuti, Batwa, Ethiopian Ancient_SouthAfrica_HG, Juhoan, Yoruba, Berom as donor populations to paint all recipient haplotypes. wRHG includes 15 Baka and Biaka individuals that shares the lowest haplotype copying probabilities with Yoruba population based on the pairwise coancestry matrix. The output copy vectors were used as input for SOURCEFINDv2 (*37*). We included various configurations by additionally including different combinations of West-Central African surrogate populations to model Bantu-related ancestry (Table S3). Each SOURCEFIND run was performed 100 independent times, each for 2 million iterations and sampling every 5,000 iterations after discarding the first 50,000 as burn-in with a maximum of 8 reference clusters allowed per target and a prior expectation of 6 reference clusters (self.copy.ind:0, num.surrogates:8, exp.num.surrogates:6). For each target, the sample with the highest posterior probability from each run was retained, and final ancestry proportions were computed as posterior-weighted averages across these retained samples. We then ran fastGLOBETROTTER (*44*) using the same copy vector and painting sample files. To date admixture events associated with the Bantu expansion, we used Non-Bantu Southern Bantoid speakers (Tikar, Bamileke, and Mundani) as the Bantu-related surrogate population. We performed 1,000 bootstrap replicates to estimate 95% confidence intervals of admixture dates for each target under both single-event and two-event models. When more than 5% of bootstrap replicates produced a two-date model with a ‘maxScore.2events’ ≥ 0.15, evidence for a two-pulse admixture model was considered and reported. For groups with fewer than four samples, we applied jackknife resampling to calculate confidence intervals by sequentially dropping one of the 22 autosomes at a time, as described in Wangkumhang et al. (2022).

### Local ancestry inference with FLARE

Local ancestry was inferred using FLARE v0.5.1 (*33*) to estimate ancestry tracts along each haploid genome of admixed Bantu-speaking individuals. Three reference panels representing the major ancestral sources inferred by fastGLOBETROTTER were constructed (West, East, and South). The West panel included western rainforest hunter-gatherers (wRHG) and non-Bantu Bantoid speakers (Bantoid); the East panel included Ethiopian, Dinka, Mbuti, and Bantoid populations; and the South panel included Ju’hoan (North and South) together with Bantoid populations. Bantu-speaking populations from these regions were analyzed as targets according to their geographic location. FLARE did not detect non-Bantu ancestry in Zambian, Mozambican, and Angolan Bantu-speaking populations, likely due to the very low proportion of such ancestry in these groups.

### Ancestry-specific genome-wide genealogy inference with Relate/CoalRate and Fitting Isolation-Migration Model

Phased VCFs from were converted to .haps and .sample formats using the ConvertFromVcf mode implemented in RelateFileFormats (*32*). Input data were then processed with PrepareInputFiles.sh from Relate, which incorporated ancestral allele information from the GRCh37_e59 human ancestor reference, applied the 1000 Genomes pilot accessibility mask while correcting inter-marker distances to account for masked regions, and annotated population labels to terminal branches in the resulting trees. Analyses were performed on the inferred trees assuming a mutation rate of 1.25 × 10⁻⁸ per generation and a haploid effective population size of 30,000, using recombination maps from the 1000 Genomes Project. Tree sequences were refined through five iterations of joint estimation of effective population sizes, mutation rates, and branch lengths using Relate’s EstimatePopulationSize.sh, with a generation time of 30 years and the parameter --bins 2,7,0.2. To improve local coalescence-rate estimations we followed two strategies for sub-setting the refined global tree: (i) each population-specific tree was separately extracted for within-population coalescence rate inference; (ii) Bantu-speaking populations and West African trees were extracted for both within- and across-population coalescence rate inference. The subtrees from (i) and (ii) were then further refined through five iterations of joint estimation of effective population sizes, mutation rates, and branch lengths using EstimatePopulationSize.sh.

We then used CoalRate (https://github.com/leospeidel/CoalRate) to estimate ancestry-specific within- and cross-population coalescence rates from the refined per-population and Bantu/West African genealogical subtrees. Tree tips of admixed Bantu individuals were masked if assigned to non-Bantu-related ancestries or labeled as uncertain when posterior ancestry probabilities were below 0.90 as estimated by FLARE. The output files containing ancestry-specific per-population coalescent rates were parsed using the relater R package (https://github.com/leospeidel/relater).

The mean and the 10th and 90th bootstrap percentiles of per-epoch coalescent rates were summarized for within-population coalescence rates to estimate effective population size trajectories over time (1/(2 × coalescent rate)). Effective population size (Ne) trajectories inferred with Relate were restricted to populations with ≥12 individuals, as smaller sample sizes produced unstable estimates.

To account for post-split migrations between Bantu speaking populations, we obtained time-dependent migration rates m(t) and cumulative migration probabilities *M*(*t*) using MSMC-IM (*35*), which fits an Isolation-with-Migration (IM) model to the distribution of ancestry-specific coalescent rates as described in (*31*). For our split time estimations, we allowed Bantu Kenya and South Africa as additional populations with slightly smaller sample sizes to improve regional representation. Although these populations produced noisy *N*_e_ trajectories, their *M*(*t*) profiles remained stable.

### Genetic Data Simulation

Genetic data were simulated using msprime v1.3.4 (*66*) under two demographic models informed by empirical genetic and archaeological data. The effective population size was initially constant at 5,000 between 100 and 50 generations ago, followed by an expansion to 150,000 over 50 generations. This was followed by a bottleneck to 10,000 over 35 generations and a subsequent expansion to 100,000 over the last 15 generations. The second model included gene flow from RHG into BSP between 30 and 40 generations ago, with a migration rate of 0.0105361. RHG was modelled with a reduction in effective population size from 10,000 to 2,000 and a divergence from BSP 4,500 generations ago (*67*) .Chromosome-specific recombination maps were used, and mutations were simulated at a rate of (1.2 × 10^-8^) per base pair per generation across all 22 autosomes.

### Recent Effective Population Size Inference using IBD segments

Both empirical and simulated data were processed following the HapNe pipeline (https://github.com/PalamaraLab/HapNe). IBD segments were inferred from VCF files using Hap-IBD (*68*) and refined following (*69*). Segments overlapping genomic gaps in the GRCh37 reference genome were removed. Gabonese individuals with high levels of IBD sharing (mean + 3 SD) were excluded within each population to reduce the impact of background relatedness. West-central African populations were merged into larger groups based on genetic structure and language families to increase sample sizes. HapNe-IBD (*42*) was then run separately for each population analysed.

### Ancestry-masked PCA and *F*_ST_

To restrict analyses to Bantu-related ancestry, genotypes were set to missing at positions where the FLARE posterior probability of Bantu-related ancestry fell below 0.8. Each masked population was downsampled to 14 individuals, matching the smallest available sample size, and variants with an allele number (AN) below 20 were excluded. The population VCF files were then merged using BCFtools. Sites present in at least one population were intersected with an LD-pruned SNP set generated using PLINK (--indep-pairwise 200 25 0.2) from a large sub-Saharan African dataset, resulting in a final set of 870,623 SNPs. For comparability, the unmasked dataset was restricted to the same set of SNP positions, and the corresponding populations in the unmasked dataset were replaced with their masked versions. Pairwise Hudson *F*_ST_ was estimated from per-population allele counts and sample sizes following (*70*), using shared SNPs for each population pair. Because ancestry masking and variable coverage cause each population pair to be evaluated on a different SNP subset, potentially inflating *F*_ST_ for more heavily masked populations, a double-centering correction was applied to the resulting matrix (*71*). PCA was performed using smartPCA (*25*), including all individuals in one analysis and projecting masked individuals onto the PCA results obtained from the unmasked dataset (lsqproject: YES).

## Acknowledgments

We warmly thank all volunteers and communities participating in this research. We thank the HPC Core Facility of Institut Pasteur (Paris) for the management of computational resources. This study makes use of data generated by H3Africa. A full list of the investigators who contributed to the generation of the data is available from https://h3africa.org/.

## Author contributions

Conceptualization: R.O., L.Q.-M., E.P.; Data curation: O.P., J.P.-M., M.R.; Formal analysis: O.P.; Funding acquisition: L.Q.-M., M.-F.D., E.P.; Investigation: O.P., M.-F.D., D.M., J.-M.H., R.G., P.M.-D., G.d.S., R.O., L.Q.-M., E.P.; Methodology: O.P., G.L., D.L., E.P.; Resources: F.M., C.H., S.K., M.-H.P., C.P., M.H., M.S., B.C., L.B.B., J.-M.H., G.d.S., P.M.-D., R.O.; Project administration: L.Y.; Software: M.R., D.L., O.P.; Supervision: M.-F.D., R.O., E.P.; Visualization: O.P.; Writing – original draft: O.P., E.P.; Writing – review & editing: O.P., L.Q.-M., E.P., with contributions from all authors.

## Diversity, equity, ethics, and inclusion

TO BE DONE.

## Data, code, and materials availability

The whole-genome sequencing dataset generated and analyzed in this study is available from the European Genome-Phenome Archive (EGA; https://www.ebi.ac.uk/ega/), under accession code EGAS00001003602. Data access and use is restricted to academic research on human history and evolution. The HGDP (*72*) and 1000 Genomes Project (*19*) genomes were retrieved from the International Genome Sample Resource (IGSR) data portal. The SGDP genome data were retrieved from the EBI European Nucleotide Archive (accession codes PRJEB9586 and ERP010710). The genome data from Choudhury *et al.* (*15*) were requested from https://catalog.h3africa.org/ and retrieved from the EGA (accession code: EGAS00001002976). The genome data from Tallman *et al.* (*16*) were retrieved from EGA (accession code EGAD00001011992). The genome data from Fan *et al.* (*20*) were retrieved from the Database of Genotypes And Phenotypes (dbGAP) (accession code: phs003096.v1.p1). The genome data from Chak *et al.* (*21*) were retrieved from EGA (accession code: EGAS50000001991).

## List of Supplementary Materials

Materials and Methods

Figs. S1 to S10

Tables S1 to S6

MDAR Reproducibility Checklist

